# Differential sphingosine-1-phosphate receptor-1 protein expression in the dorsolateral prefrontal cortex between schizophrenia Type 1 and Type 2

**DOI:** 10.1101/2021.05.15.444302

**Authors:** Ganesh B. Chand, Hao Jiang, J. Philip Miller, C. Harker Rhodes, Zhude Tu, Dean F. Wong

## Abstract

Understanding the etiology and treatment approaches in schizophrenia is challenged in part by the heterogeneity of this disorder. One encouraging progress is the growing evidence that there are subtypes of schizophrenia that may relate to disease duration and premorbid severity. Recent *in vitro* findings of messenger ribonucleic acid (mRNA) gene expression on postmortem dorsolateral prefrontal cortex (DLPFC) showed that schizophrenia has two subtypes, those with a relatively normal DLPFC transcriptome (Type 1) and those with differentially expressed genes (Type 2). Sphingosine-1-phosphate receptor-1 (S1PR1) is one of the genes that was highly upregulated in Type 2 compared to Type 1 and controls. The impact of that finding is limited because it only can be confirmed through analysis of autopsy tissue, and the clinical characteristics such as symptoms severity or illness duration was not available from that Medical Examiner based autopsy study. However, S1PR1 has great potential because it is a target gene that can be accessed via positron emission tomography (PET) *in vivo* using specific radioligands (starting with [^11^C]CS1P1) successfully developed at our center in human brain imaging. As a preliminary study to validate this PET target in schizophrenia, S1PR1 protein expression was assessed by receptor autoradiography (ARG) using [^3^H]CS1P1 and immunohistochemistry (IHC) in the DLPFC from patients with schizophrenia classified as Type 1 or Type 2 based on their DLPFC transcriptomes and from controls. Our analyses demonstrate that ARG S1PR1 protein expression is significantly higher in Type 2 compared to Type 1 (p < 0.05) and controls (p < 0.05), which was consistent with previous mRNA S1PR1. These findings support the possibility that PET S1PR1 can be used as a future imaging biomarker to distinguish these subgroups of schizophrenic patients during life with obvious implications for both patient management and the design of clinical trials to validate novel pharmacologic therapies.

## Introduction

Schizophrenia is a neuropsychiatric condition that currently affects ∼3 million people in the United States and ∼7.8 billion people worldwide (Chong et al., 2016; Cloutier et al., 2016; McCutcheon et al., 2020). Individuals with schizophrenia exhibit highly heterogeneous genetic profiles (Arnedo et al., 2015), clinical symptoms (Derks et al., 2012), illness course (Carpenter and Kirkpatrick, 1988; Huber, 1997), treatment response (Malhotra, 2015; Palaniyappan et al., 2013), and neuroimaging markers (Nenadic et al., 2015; Voineskos et al., 2013; Voineskos et al., 2020). Despite extensive efforts, understanding schizophrenia mechanisms and developing targeted personalized clinical treatment are not encouraging (Insel and Cuthbert, 2015; Kapur et al., 2012). Recent findings by Bowen and colleagues (Bowen et al., 2019) suggest that the mRNA expression can be used to divide schizophrenic patients into two types, Type 1 schizophrenia patients with an essentially normal transcriptome in their dorsolateral prefrontal cortex (DLPFC) and Type 2 schizophrenia patients with hundreds of differentially expressed genes in their DLPFC.

A serious limitation of mRNA expression studies like that of Bowen and colleagues (Bowen et al., 2019) is that they require brain tissue which is generally not available except at autopsy. Fortunately PET ligands for one of the genes differentially expressed in the Type 2 schizophrenic brains, sphingosine-1-phosphate receptor-1 (S1PR1), have been recently developed at Washington University School of Medicine (Jiang et al., 2021; Jin et al., 2017; Liu et al., 2017; Liu et al., 2016; Liu et al., 2020; Luo et al., 2019). S1PR1 radioligand has gained significant interest for *in vivo* targeted imaging of inflammation in brain diseases, with the recent FDA-approved S1PR1-based treatments, such as Fingolimod, Siponimod, and Ozanimod (Bross et al., 2020) for multiple sclerosis. The present study examines the differential expression of S1PR1 in the DLPFC of Type 1 and Type 2 schizophrenic patients at the protein level as a preliminary step towards the use of PET to distinguish Type 1 from Type 2 schizophrenia during life. We performed autoradiography (ARG) and immunohistochemistry (IHC) analyses in DLPFC tissues of controls, Type 1 and Type 2 schizophrenic patients. We hypothesized that S1PR1 protein expression will show elevations in Type 2 schizophrenia compared to Type 1 schizophrenia and controls consistent with Bowen’s S1PR1 mRNA findings (Bowen et al., 2019).

## Materials and methods

### Human brain tissues

Human brain tissues were obtained from the Human Brain Collection Core (HBCC) at the National Institute of Mental Health (NIMH). These tissues corresponded to the same tissues studied by Bowen and colleagues (Bowen et al., 2019) because any other tissues from different sources would be difficult to distinguish Type 1 and Type 2 schizophrenia. Tissues were used in accordance with the guidelines of Washington University in St. Louis. All samples were stored at -80 ºC at the Washington University’s Radiology labs until used.

### *In vitro* immunohistochemistry staining study

*In vitro* IHC staining of S1PR1 was carried out in frozen sections from human DLPFC. All sections were pre-warmed at room temperature (RT) for 5 minutes and then fixed with 4% paraformaldehyde in phosphate buffered saline (PBS) for 10 minutes at RT, washed 3 times in PBS, and then heated in boiling water bath in antigen retrieval buffer for 30 minutes. Sections were then rinsed with PBS and blocked with 5% horse serum for 1 hour at RT. After that, all sections were stained with anti-S1PR1 antibody (Alomone, Jerusalem, Israel) overnight at 4ºC, washed and followed by incubation with ImmPRESS HRP Horse anti-rabbit polymer for 1 hour at RT, and developed using ImmPACT DAB (Vector Laboratories, Burlingame, CA). Hematoxylin and eosin (H&E) staining was also performed in adjacent slides to identify gray and white matters in the brain.

### *In vitro* autoradiography study

*In vitro* ARG study was carried out in frozen sections from human DLPFC using [^3^H]CS1P1 for S1PR1 receptor protein. Sections were pre-warmed to RT, and then incubated with Hank’s balanced salt solution (HBSS) buffer containing 10 mM HEPES, 5mM MgCl_2_, 0.2% BSA, and 0.1mM EDTA at pH7.4 for 5 minutes at RT in a coplin jar. All sections were then incubated with 0.5nM [^3^H]CS1P1 for 30 min in a coplin jar with gentle shaking at RT. After that, all sections were washed with a buffer for 3 minutes for three times, and then rinsed in ice-cold H_2_O for 1 min and air dried overnight. Slides were incubated with Carestream BioMax Maximum Sensitivity ARG film (Carestream, Rochester, NY) in a Hypercassette ARG cassette (Cytiva, Amersham, UK) for 30 days along with an ART-123 Tritium Standards (American Radiolabeled Chemicals, St Louis, MO). The film was processed using a Kodak film developer (Kodak, Rochester, NY). To determine the non-specific binding, 10 μM of S1PR1 specific antagonist NIBR-0213 (Cayman, Ann Arbor, MI) was introduced and incubated with samples as described above. The image was processed and analyzed using Fiji ImageJ. Brain regions of interest (ROIs) were selected from the ARG images according to the hematoxylin staining in the adjacent slide. ROIs were randomly selected from different regions of the DLPFC gray matter, and the intensity was measured and calculated in fmol/mg.

### Statistical analysis

In statistical analysis we fitted a mixed repeated measure model to take into account the triplicate measure variability (SD_R_) [with a compound symmetry covariance structure] as well as the interpersonal variability (within group) SD_S_. In this statistical analysis, we modeled that the individual measures had a variance of SD_R_^2^ + SD_S_^2^. As this involved different groups of subjects, the appropriate variance for the group test was chosen as SD_S_^2^. The F-test for the null that all three groups have the same mean was tested. To compare between groups, a one-tailed F-test was used, justified by the prior work (Bowen et al., 2019). A p-value ≤ 0.05 was considered statistically significant.

## Results

### Postmortem human subjects

DLPFC tissues from twenty human subjects including ten neurologically normal controls, five Type 1 schizophrenia subjects, and five Type 2 schizophrenia subjects were used in this study (**Table 1**). Mean age (standard deviation) was 55.20 years (9.69) of normal controls, 56.40 years (8.32) of Type 1 schizophrenia, and 53.20 years (10.26) of Type 2 schizophrenia subjects. There was one female in controls, none in Type 1 schizophrenia, and one in Type 2 schizophrenia. There were five Caucasians and five African-Americans in controls, three Caucasians and two African-Americans in Type 1 schizophrenia, and two Caucasians and three African-Americans in Type 2 schizophrenia. Age, sex, and race were not significantly different among groups.

**Table 1:**
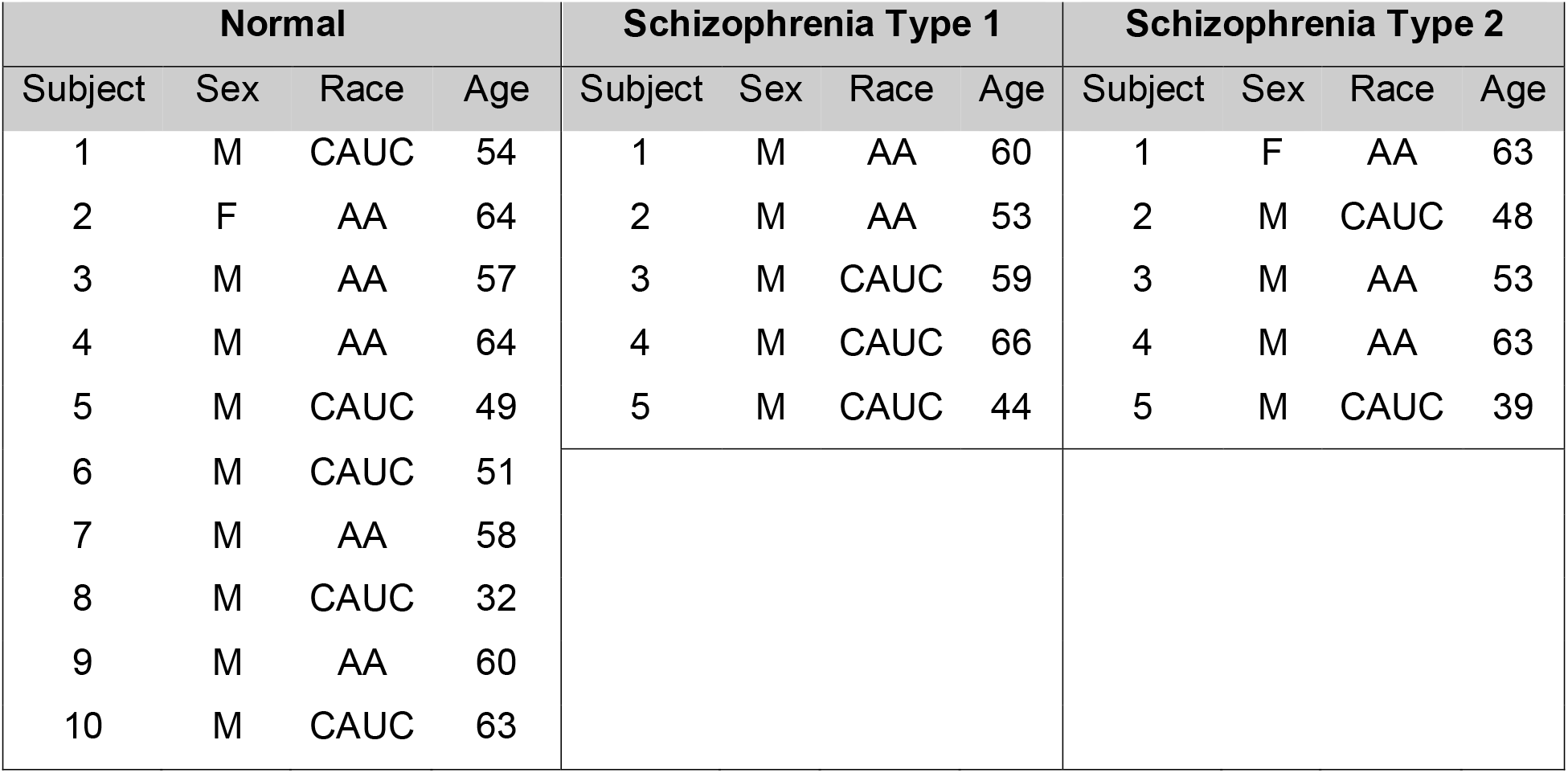
Human DLPFC tissues used in this study from normal controls, schizophrenia Type 1 and Type 2 patients. (F: Female; M: Male; AA: African-American; CAUC: Caucasian; Age in years).

| Normal |  |  |  | Schizophrenia Type 1 |  |  |  | Schizophrenia Type 2 |  |  |  |
| --- | --- | --- | --- | --- | --- | --- | --- | --- | --- | --- | --- |
| Subject | Sex | Race | Age | Subject | Sex | Race | Age | Subject | Sex | Race | Age |
| 1 | M | CAUC | 54 | 1 | M | AA | 60 | 1 | F | AA | 63 |
| 2 | F | AA | 64 | 2 | M | AA | 53 | 2 | M | CAUC | 48 |
| 3 | M | AA | 57 | 3 | M | CAUC | 59 | 3 | M | AA | 53 |
| 4 | M | AA | 64 | 4 | M | CAUC | 66 | 4 | M | AA | 63 |
| 5 | M | CAUC | 49 | 5 | M | CAUC | 44 | 5 | M | CAUC | 39 |
| 6 | M | CAUC | 51 |  |  |  |  |  |  |  |  |
| 7 | M | AA | 58 |  |  |  |  |  |  |  |  |
| 8 | M | CAUC | 32 |  |  |  |  |  |  |  |  |
| 9 | M | AA | 60 |  |  |  |  |  |  |  |  |
| 10 | M | CAUC | 63 |  |  |  |  |  |  |  |  |

### Immunostaining of S1PR1

Immunostaining of S1PR1 was performed in control and schizophrenia samples (**Figures 1 and 2**). In general, S1PR1 was mainly expressed in the gray matter of the DLPFC. In particularly, the expression of S1PR1 was relatively high in outer granular layer, outer pyramidal layer, inner granular layer, inner pyramidal layer, and multiform layer with very low to no amount in the molecular layer of gray matter and white matter. No significant morphological difference was identified between normal control and schizophrenia subjects.

**Figure 1:**
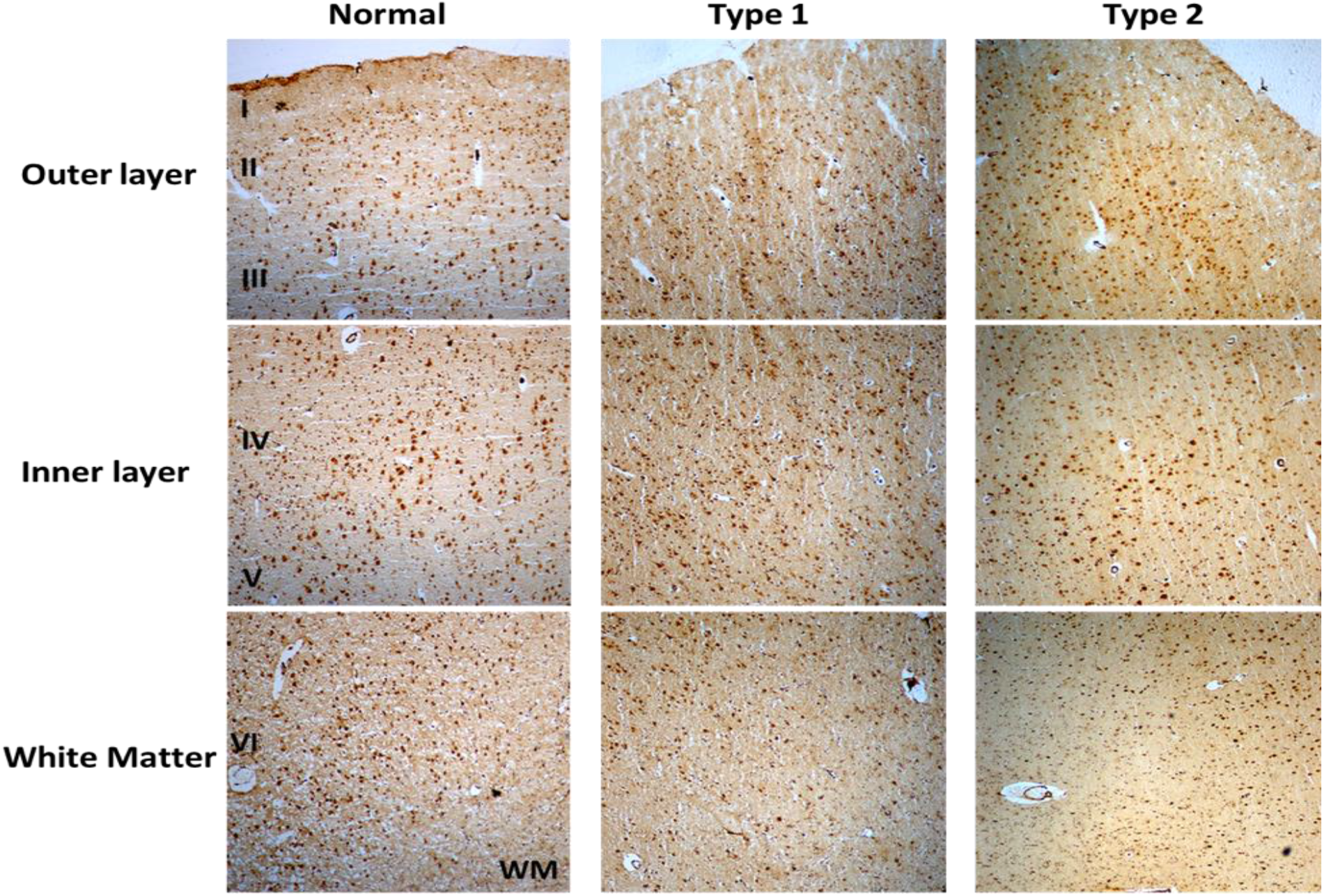
Immunohistochemistry (IHC) of S1PR1 in postmortem DLPFC tissues from the representative normal control and schizophrenia Type 1 and Type 2.

**Figure 2:**
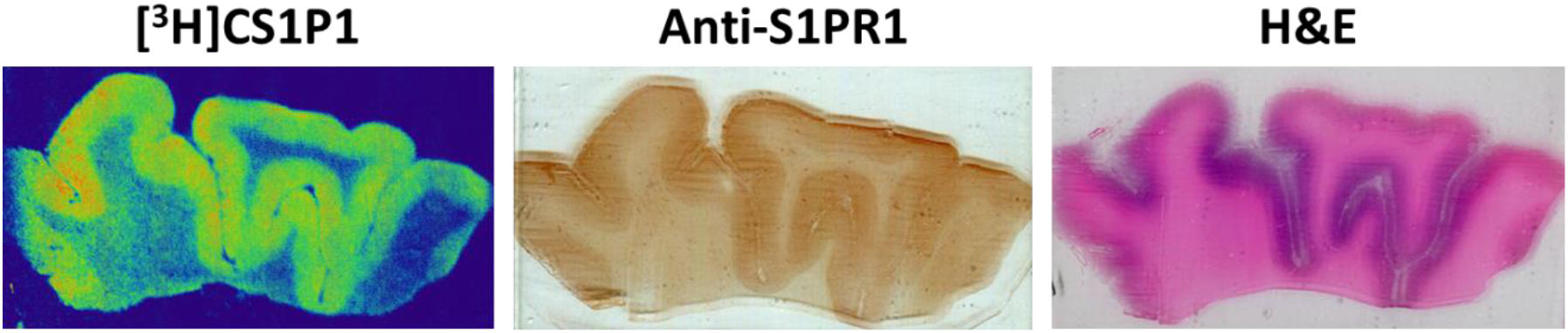
Representative images of [^3^H]CS1P1 autoradiograph, S1PR1 immunostaining, and Hematoxylin and eosin (H&E) staining in postmortem human DLPFC tissues. The distribution of [^3^H]CS1P1 matched well with anti-S1PR1 antibody, and was mainly located in the gray matter regions as indicated in the H&E staining.

### Autoradiography of S1PR1 specific [^3^H]CS1P1

ARG analysis of S1PR1 was performed in control and schizophrenia DLPFC samples.

The distribution pattern of [^3^H]CS1P1 matched well with immunostaining analysis using S1PR1 specific antibody, indicating [^3^H]CS1P1 is specific to S1PR1 in postmortem human tissues (**Figures 2 and 3**). Similar to S1PR1 immunostaining analysis, S1PR1 specific [^3^H]CS1P1 was mainly distributed in the gray matter of DLPFC, with no to very low amount of [^3^H]CS1P1 distributed in the white matter region. In addition, blocking study using S1PR1 antagonist NIBR-0213 showed significant reduction of [^3^H]CS1P1 indicating the [^3^H]CS1P1 is specific to S1PR1 (**Figure 3**). Compared with IHC analysis, ARG provides both quantification and localization of radioligand at the same time in distinct anatomical structures, and enables us to quantify the absolute amount of [^3^H]CS1P1 in control and different types of schizophrenia subjects.

**Figure 3:**
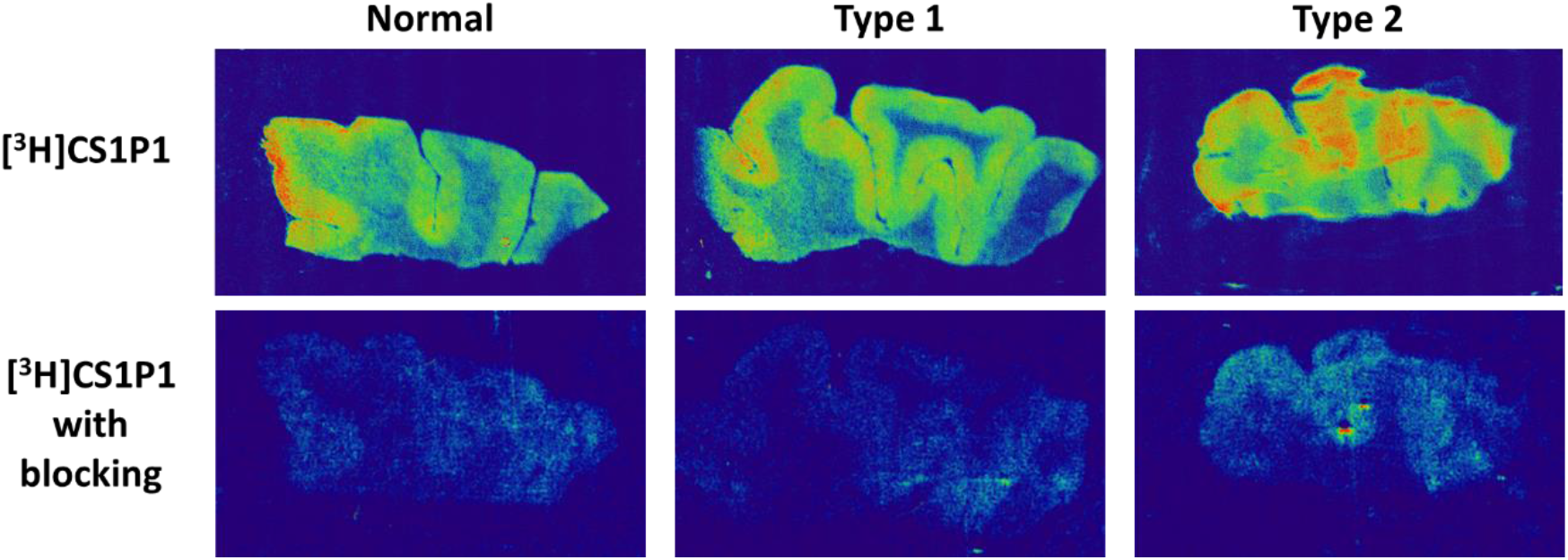
Autoradiography images of S1PR1 using [^3^H]CS1P1 in postmortem DLPFC tissues from representative normal control, schizophrenia Type 1, and schizophrenia Type 2. In general, [^3^H]CS1P1 was higher in Type 2 schizophrenia subjects compared with normal control and Type 1 schizophrenia subjects.

ARG S1PR1 intensity in the DLPFC was compared between the groups. In general, the intensity of [^3^H]CS1P1 was higher in Type 2 schizophrenia subjects compared with normal control and Type 1 schizophrenia subjects (**Figure 3**). For statistical comparison of results, SD_R_ and SD_s_ were estimated. The estimates were SD_R_ = 6.81 and SD_S_ = 12.52. The F-test for the null that all three groups have the same mean was F(2,17) = 3.49, p = 0.05. A one-tailed F-test for controls vs. Type 2 was p = 0.021, Type 2 vs. Type 1 was p = 0.012, and controls vs. average of Type 1 and Type 2 was p = 0.18 (**Table 2**).

**Table 2:** Summary of AGR S1PR1 intensity (in fmol/mg) in the DLPFC from normal control, schizophrenia Type 1, and schizophrenia Type 2 groups. (SE: standard error of mean).

| **Group** | **N** | **ARG S1PR1**<br>**(mean)** | **ARG S1PR1**<br>**(SE)** |
| --- | --- | --- | --- |
| **Normal** | 10 | 76.10 | 4.15 |
| **Type 1** | 5 | 71.44 | 5.87 |
| **Type 2** | 5 | 91.91 | 5.87 |

## Discussion

In this study, we evaluated the expression of S1PR1 and distribution of S1PR1 specific tracer [^3^H]CS1P1 in human DLPFC tissues from normal control, Type 1 and Type 2 schizophrenia subjects. Our data showed, in the DLPFC of humans, S1PR1 is highly expressed in the gray matter region with much lower expression in the white matter regions. Similar to the immunostaining study, ARG analysis using [^3^H]CS1P1 showed a relatively high tracer uptake in the gray matter of DLPFC whereas no to very low amount of [^3^H]CS1P1 was identified in the white matter region. Tracer uptake in the Type 2 schizophrenia samples was significantly higher than the Type 1 schizophrenia samples and normal controls.

The present ARG findings are consistent with a previous study in other regions of the human brain (Nishimura et al., 2010) that suggests S1PR1 is mainly localized in gray matter and confirmed the specificity of our S1PR1 specific radioligand [^3^H]CS1P1. This study extends the previous findings (Bowen et al., 2019) by demonstrating that S1PR1 protein as well as mRNA is differentially expressed in the DLPFC of Type 2 schizophrenic patients. The trend of tracer uptake between control and all schizophrenia subjects was similar to previous findings using RT-PCR (Esaki et al., 2020). For the two types of schizophrenia patients, it appeared that only Type 2 schizophrenia has significantly upregulated S1PR1 compared to Type 1 schizophrenia and controls, respectively. The fact that S1PR1 IHC did not distinguish Type 1 from Type 2 patients was not unexpected. IHC is a highly non-linear, at best semi-quantitative technique with excellent spatial resolution, but limited ability to detect quantitative changes in gene expression. The ∼28% increase in S1PR1 expression in Type 2 as compared to Type 1 patients is not expected to be detectable by IHC. Thus, the present ARG findings and previous reports taken together indicate that both S1PR1 protein and S1PR1 mRNA upregulate in DLPFC only in a subset of schizophrenia.

These results have important implications for the future schizophrenia subtype-based studies with *in vivo* PET [^11^C]CS1P1. The [^11^C]CS1P1 has recently emerged as a promising radiotracer for *in vivo* PET imaging of neuro-inflammation (Liu et al., 2017; Liu et al., 2016; Liu et al., 2020; Luo et al., 2019). Neuro-inflammation has been associated with schizophrenia throughout the literature (Comer et al., 2020; Najjar and Pearlman, 2015). This has included elevated cytokines (Potvin et al., 2008; Upthegrove et al., 2014) that may be due to activated microglia. The neuro-inflammatory observation seems to be most associated with early illness stages and patients with acute psychosis symptom exacerbations (Kraguljac et al., 2021). The current standard in Translocator protein (18 kDa) (TSPO) PET studies for investigating neuro-inflammation in schizophrenia shows discrepancies in outcomes (Marques et al., 2019). Moreover, this method requires the additional step and cost of a priori genotyping and most seriously the reduction of sample size by 1/3 or 2/3 depending on whether medium and high affinity binding or high only subjects are ultimately used in the analysis. Studies with TSPO PET 1^st^ and 2^nd^ generation radioligands have not convincingly shown elevations in chronic schizophrenia (Kenk et al., 2015; Takano et al., 2010). TSPO suffers from variable signal to noise and, more seriously, the need to genotype subjects where high and possibly mid-affinity binders can be studied but omitting those with low-affinity binders. Recent reviews have cast doubt on TSPO as a schizophrenia imaging target (Sneeboer et al., 2020), and meta-analyses have shown only small elevations and others none (Marques et al., 2019). Recent reviews have even suggested decreases in schizophrenia and acute psychosis (Meyer et al., 2020). Despite inconsistency with TSPO PET, studies have shown elevations of plasma inflammatory markers, including IL-6 in plasma and CSF (Coughlin et al., 2016). Cox-2 is elevated in schizophrenia, but PET ligands for inflammation response are not yet well developed (Prabhakaran et al., 2021), nor for cannabinoid (CB2) (Banaszkiewicz et al., 2020). Thus there is great potential for studying neuro-inflammation (or related mechanisms) with S1PR1 PET ligands to fill an important gap in the theory of neuro-inflammation in schizophrenia to the realization of *in vivo* measures. Modulators targeting S1PR1 are already FDA-approved therapeutics of treating multiple sclerosis (Bross et al., 2020; Marciniak et al., 2020), which could further accelerate its important role for treatment development in schizophrenia.

One limitation is that S1PR1 mRNA study (Bowen et al., 2019) can’t be used to identify schizophrenia Type 1 and Type 2 *in vivo* without PET and S1PR1 specific radiotracer. However, there is mounting evidence that schizophrenia has two neuroanatomical types *in vivo* (Chand et al., 2020). This recent study by Chand and colleagues (Chand et al., 2020) thus can be used to divide schizophrenia patients into types *in vivo* using structural MRI so that PET S1PR1 can be evaluated subsequently in schizophrenia types. We also acknowledge that the sample size is relatively small for this study, and the future studies should focus on replicating these findings in larger samples. However, we were limited by the available tissues in the HBCC NIMH brain bank that corresponded to the same tissues studied by Bowen and colleagues (Bowen et al., 2019), and any other tissues would be difficult to distinguish Type 1 and Type 2 schizophrenia. In addition, it remains to be investigated S1PR1 protein and S1PR1 mRNA expressions in other brain regions besides DLPFC of controls and Type 1 and Type 2 schizophrenia patients.

In conclusion, the present study evaluated DLPFC postmortem tissues from controls, schizophrenia Type 1 and Type 2, demonstrated S1PR1 protein is highly expressed in gray matter region, and most importantly showed that only Type 2 schizophrenia has upregulated S1PR1 protein expression in line with previous S1PR1 mRNA results. Overall, these findings strongly suggest S1PR1 might serve as a candidate target in schizophrenia subtypes with PET where protein is the target.

## Acknowledgement

Authors would like to acknowledge the Human Brain Collection Core (HBCC), National Institute of Mental Health (NIMH) for human DLPFC postmortem tissues, S. Marenco, MD, Acting Director of HBCC, and Mallinckrodt Institute of Radiology internal funds (D.F.W.).

## Data availability statement

Human brain tissue data used in this study are publicly available from the Human Brain Collection Core (HBCC) at the National Institute of Mental Health (NIMH) following the data request procedure. The original contributions presented in the study are included in the article, and further inquiries can be directed to the corresponding author (G.B.C.).

## References

Arnedo, J., Svrakic, D.M., Del Val, C., Romero-Zaliz, R., Hernandez-Cuervo, H., Molecular Genetics of Schizophrenia, C., Fanous, A.H., Pato, M.T., Pato, C.N., de Erausquin, G.A., Cloninger, C.R., Zwir, I., 2015. Uncovering the hidden risk architecture of the schizophrenias: confirmation in three independent genome-wide association studies. Am J Psychiatry 172, 139–153.

Banaszkiewicz, I., Biala, G., Kruk-Slomka, M., 2020. Contribution of CB2 receptors in schizophrenia-related symptoms in various animal models: Short review. Neurosci Biobehav Rev 114, 158–171.

Bowen, E.F.W., Burgess, J.L., Granger, R., Kleinman, J.E., Rhodes, C.H., 2019. DLPFC transcriptome defines two molecular subtypes of schizophrenia. Transl Psychiatry 9, 147.

Bross, M., Hackett, M., Bernitsas, E., 2020. Approved and Emerging Disease Modifying Therapies on Neurodegeneration in Multiple Sclerosis. Int J Mol Sci 21.

Carpenter, W.T., Kirkpatrick, B., 1988. The heterogeneity of the long-term course of schizophrenia. Schizophr Bull 14, 645–652.

Chand, G.B., Dwyer, D.B., Erus, G., Sotiras, A., Varol, E., Srinivasan, D., Doshi, J., Pomponio, R., Pigoni, A., Dazzan, P., Kahn, R.S., Schnack, H.G., Zanetti, M.V., Meisenzahl, E., Busatto, G.F., Crespo-Facorro, B., Pantelis, C., Wood, S.J., Zhuo, C., Shinohara, R.T., Shou, H., Fan, Y., Gur, R.C., Gur, R.E., Satterthwaite, T.D., Koutsouleris, N., Wolf, D.H., Davatzikos, C., 2020. Two distinct neuroanatomical subtypes of schizophrenia revealed using machine learning. Brain 143, 1027–1038.

Chong, H.Y., Teoh, S.L., Wu, D.B.C., Kotirum, S., Chiou, C.F., Chaiyakunapruk, N., 2016. Global economic burden of schizophrenia: a systematic review. Neuropsychiatric Disease and Treatment 12 357–373.

Cloutier, M., Aigbogun, M.S., Guerin, A., Nitulescu, R., Ramanakumar, A.V., Kamat, S.A., DeLucia, M., Duffy, R., Legacy, S.N., Henderson, C., Francois, C., Wu, E., 2016. The Economic Burden of Schizophrenia in the United States in 2013. J Clin Psychiatry 77, 764–771.

Comer, A.L., Carrier, M., Tremblay, M.E., Cruz-Martin, A., 2020. The Inflamed Brain in Schizophrenia: The Convergence of Genetic and Environmental Risk Factors That Lead to Uncontrolled Neuroinflammation. Front Cell Neurosci 14, 274.

Coughlin, J.M., Wang, Y., Ambinder, E.B., Ward, R.E., Minn, I., Vranesic, M., Kim, P.K., Ford, C.N., Higgs, C., Hayes, L.N., Schretlen, D.J., Dannals, R.F., Kassiou, M., Sawa, A., Pomper, M.G., 2016. In vivo markers of inflammatory response in recent-onset schizophrenia: a combined study using [(11)C]DPA-713 PET and analysis of CSF and plasma. Transl Psychiatry 6, e777.

Derks, E.M., Allardyce, J., Boks, M.P., Vermunt, J.K., Hijman, R., Ophoff, R.A., Group, 2012. Kraepelin was right: a latent class analysis of symptom dimensions in patients and controls. Schizophr Bull 38, 495–505.

Esaki, K., Balan, S., Iwayama, Y., Shimamoto-Mitsuyama, C., Hirabayashi, Y., Dean, B., Yoshikawa, T., 2020. Evidence for Altered Metabolism of Sphingosine-1-Phosphate in the Corpus Callosum of Patients with Schizophrenia. Schizophr Bull 46, 1172–1181.

Huber, G., 1997. The heterogeneous course of schizophrenia. Schizophr Res 28, 177–185.

Insel, T.R., Cuthbert, B.N., 2015. Brain disorders? Precisely. Science 348 (6234), 499–500.

Jiang, H., Joshi, S., Liu, H., Mansor, S., Qiu, L., Zhao, H., Whitehead, T., Gropler, R.J., Wu, G.F., Cross, A.H., Benzinger, T.L.S., Shoghi, K.I., Perlmutter, J.S., Tu, Z., 2021. In Vitro and In Vivo Investigation of S1PR1 Expression in the Central Nervous System Using [(3)H]CS1P1 and [(11)C]CS1P1. ACS Chem Neurosci 12, 3733–3744.

Jin, H., Yang, H., Liu, H., Zhang, Y., Zhang, X., Rosenberg, A.J., Liu, Y., Lapi, S.E., Tu, Z., 2017. A promising carbon-11-labeled sphingosine-1-phosphate receptor 1-specific PET tracer for imaging vascular injury. J Nucl Cardiol 24, 558–570.

Kapur, S., Phillips, A.G., Insel, T.R., 2012. Why has it taken so long for biological psychiatry to develop clinical tests and what to do about it? Mol Psychiatry 17, 1174–1179.

Kenk, M., Selvanathan, T., Rao, N., Suridjan, I., Rusjan, P., Remington, G., Meyer, J.H., Wilson, A.A., Houle, S., Mizrahi, R., 2015. Imaging neuroinflammation in gray and white matter in schizophrenia: an in-vivo PET study with [18F]-FEPPA. Schizophr Bull 41, 85–93.

Kraguljac, N.V., McDonald, W.M., Widge, A.S., Rodriguez, C.I., Tohen, M., Nemeroff, C.B., 2021. Neuroimaging Biomarkers in Schizophrenia. Am J Psychiatry 178, 509–521.

Liu, H., Jin, H., Yue, X., Han, J., Baum, P., Abendschein, D.R., Tu, Z., 2017. PET Study of Sphingosine-1-Phosphate Receptor 1 Expression in Response to Vascular Inflammation in a Rat Model of Carotid Injury. Mol Imaging 16, 1536012116689770.

Liu, H., Jin, H., Yue, X., Luo, Z., Liu, C., Rosenberg, A.J., Tu, Z., 2016. PET Imaging Study of S1PR1 Expression in a Rat Model of Multiple Sclerosis. Mol Imaging Biol 18, 724–732.

Liu, H., Luo, Z., Gu, J., Jiang, H., Joshi, S., Shoghi, K.I., Zhou, Y., Gropler, R.J., Benzinger, T.L.S., Tu, Z., 2020. In vivo Characterization of Four (18)F-Labeled S1PR1 Tracers for Neuroinflammation. Mol Imaging Biol 22(5), 1362–1369.

Luo, Z., Gu, J., Dennett, R.C., Gaehle, G.G., Perlmutter, J.S., Chen, D.L., Benzinger, T.L.S., Tu, Z., 2019. Automated production of a sphingosine-1 phosphate receptor 1 (S1P1) PET radiopharmaceutical [(11)C]CS1P1 for human use. Appl Radiat Isot 152, 30–36.

Malhotra, A.K., 2015. Dissecting the Heterogeneity of Treatment Response in First-Episode Schizophrenia. Schizophr Bull 41, 1224–1226.

Marciniak, A., Camp, S.M., Garcia, J.G.N., Polt, R., 2020. In silico Docking Studies of Fingolimod and S1P1 Agonists. Front Pharmacol 11, 247.

Marques, T.R., Ashok, A.H., Pillinger, T., Veronese, M., Turkheimer, F.E., Dazzan, P., Sommer, I.E.C., Howes, O.D., 2019. Neuroinflammation in schizophrenia: meta-analysis of in vivo microglial imaging studies. Psychol Med 49, 2186–2196.

McCutcheon, R.A., Reis Marques, T., Howes, O.D., 2020. Schizophrenia-An Overview. JAMA Psychiatry 77, 201–210.

Meyer, J.H., Cervenka, S., Kim, M.-J., Kreisl, W.C., Henter, I.D., Innis, R.B., 2020. Neuroinflammation in psychiatric disorders: PET imaging and promising new targets. The Lancet Psychiatry 7, 1064–1074.

Najjar, S., Pearlman, D.M., 2015. Neuroinflammation and white matter pathology in schizophrenia: systematic review. Schizophr Res 161, 102–112.

Nenadic, I., Yotter, R.A., Sauer, H., Gaser, C., 2015. Patterns of cortical thinning in different subgroups of schizophrenia. Br J Psychiatry 206, 479–483.

Nishimura, H., Akiyama, T., Irei, I., Hamazaki, S., Sadahira, Y., 2010. Cellular localization of sphingosine-1-phosphate receptor 1 expression in the human central nervous system. J Histochem Cytochem 58, 847–856.

Palaniyappan, L., Marques, T.R., Taylor, H., Handley, R., Mondelli, V., Bonaccorso, S., Giordano, A., McQueen, G., DiForti, M., Simmons, A., David, A.S., Pariante, C.M., Murray, R.M., Dazzan, P., 2013. Cortical folding defects as markers of poor treatment response in first-episode psychosis. JAMA Psychiatry 70, 1031–1040.

Potvin, S., Stip, E., Sepehry, A.A., Gendron, A., Bah, R., Kouassi, E., 2008. Inflammatory cytokine alterations in schizophrenia: a systematic quantitative review. Biol Psychiatry 63, 801–808.

Prabhakaran, J., Molotkov, A., Mintz, A., Mann, J.J., 2021. Progress in PET Imaging of Neuroinflammation Targeting COX-2 Enzyme. Molecules 26.

Sneeboer, M.A.M., van der Doef, T., Litjens, M., Psy, N.B.B., Melief, J., Hol, E.M., Kahn, R.S., de Witte, L.D., 2020. Microglial activation in schizophrenia: Is translocator 18kDa protein (TSPO) the right marker? Schizophr Res 215, 167–172.

Takano, A., Arakawa, R., Ito, H., Tateno, A., Takahashi, H., Matsumoto, R., Okubo, Y., Suhara, T., 2010. Peripheral benzodiazepine receptors in patients with chronic schizophrenia: a PET study with [11C]DAA1106. Int J Neuropsychopharmacol 13, 943–950.

Upthegrove, R., Manzanares-Teson, N., Barnes, N.M., 2014. Cytokine function in medication-naive first episode psychosis: a systematic review and meta-analysis. Schizophr Res 155, 101–108.

Voineskos, A.N., Foussias, G., Lerch, J., Felsky, D., Remington, G., Rajji, T.K., Lobaugh, N., Pollock, B.G., Mulsant, B.H., 2013. Neuroimaging evidence for the deficit subtype of schizophrenia. JAMA Psychiatry 70, 472–480.

Voineskos, A.N., Jacobs, G.R., Ameis, S.H., 2020. Neuroimaging Heterogeneity in Psychosis: Neurobiological Underpinnings and Opportunities for Prognostic and Therapeutic Innovation. Biol Psychiatry 88, 95–102.

